# Flow transports extracellular lipid-anchored proteins across the surface of living COS-7 cells

**DOI:** 10.1101/2025.08.27.672542

**Authors:** Sreeja Sasidharan, Leah Knepper, Larissa Socrier, Liliane Smits, Samuel Pash, Linda Lowe-Krentz, Damien Thévenin, Aurelia Honerkamp-Smith

## Abstract

The rapid diffusion of membrane lipids and membrane proteins in living cell plasma membranes demonstrates that the membrane is fluid. However, motion of membrane molecules is inhibited on one side by the cytoskeletal mesh, and on the other by the glycocalyx, a layer of proteoglycans with long polysaccharide chains that covers the membrane surface. A variety of biological fluid flows (including blood circulation, cilia-driven flows, and swimming motion of microorganisms) apply shear stress to cell surfaces. Cell responses to these flows govern important physiological processes such as blood pressure and immune activation. The presence of the glycocalyx is generally thought to shield cell membranes from shear stress that arises from flow. However, here we show that two different proteins, each attached by a lipid anchor to the extracellular membrane surface of living COS-7 cells, formed reversible, cell-wide concentration gradients in the direction of applied flow. Protein redistribution occurred within minutes after we applied shear stress levels commonly found in animal cardiovascular systems. The dynamic and spatial features of these gradients were consistent with passive transport by flow. Passive flow transport could be a general mechanism for spatial organization of membrane proteins. This mechanism may explain protein patterning previously observed on flow-exposed cells, and potentially forms an initial step in flow sensing.

## Introduction

Flow sensing by individual cells regulates important physiological processes. For example, osteocytes regulate bone density by detecting flows of interstitial fluid that result from the bone deformation that occurs during movements such as walking (1). Endothelial cells line blood vessels and are subjected to shear stress from blood flow; this directs a variety of responses including cell elongation and alignment, directional migration, and secretion of nitric oxide, a blood pressure regulating signal (2–4). Unidirectional, undisturbed flow activates anti-inflammatory, anti-oxidative genes, whereas disturbed flow stimulates cell proliferation and the expression of inflammatory genes (5). When chronically activated, this flow-mediated inflammation leads to development of atherosclerosis (6, 7).

The apical membrane of an endothelial cell, and the membrane proteins located there, form a likely site for flow mechanosensing. However, the specific molecular mechanism for this process remains unknown. Efforts to identify a protein “flow sensor” on endothelial cells have been inconclusive. Such a protein sensor would need to detect the tiny forces applied by cardiovascular flow, despite the shielding of these forces by the thick layer of proteins and sugars that extends from the cell membrane.

Typical estimated shear stress on blood vessel walls ranges from 1-4 Pa in arteries, and 0.1 - 0.6 Pa in veins. In small vessels, local geometry and close proximity to individual red blood cells can result in higher shear stresses (4). In the human aorta, a typical shear stress of 1 Pa would exert a force of only 0.1 femtonewton (fN) on an area equivalent to that of a single protein, 100 nm^2^ (6). This force is not large enough to change protein configuration: known mechanosensitive proteins respond to forces measured in tens of piconewtons (8–10). Despite this, endothelial cells reliably respond to the small shear stresses that arise from blood circulation (6).

A layer of glycoproteins, proteoglycans and their associated polysaccharides, and glycosylated lipids called the glycocalyx covers the membrane of almost all mammalian cells. The endothelial cell glycocalyx is particularly well studied. Estimates for its thickness range from hundreds of nanometers to several micrometers (11); measurement is challenging since it is soft, easily degraded by common imaging methods, varies with tissue type, and is highly sensitive to culture conditions in vitro. Glycocalyx loss or degradation is associated with pathological conditions such as inflammation and sepsis (2, 12). Even a glycocalyx with nanometer-scale thickness should attenuate the already small forces from circulatory flow. Foundational theoretical work predicted that shear stress at the cell membrane is negligible due to the presence of the endothelial glycocalyx (13, 14).

On the other hand, femtonewton-sized forces can transport lipid-anchored proteins laterally across a simplified model of the cell plasma membrane, a lipid bilayer supported on glass. Flow creates hydrodynamic drag on proteins and other biomolecules that protrude from a membrane. Our previous experiments showed that streptavidin bound to biotinylated lipids was rapidly redistributed across cell-sized membrane patches under one Pa of shear stress (the corresponding force applied to an individual streptavidin protein is 0.1 fN) (15). We also found that the shape of an individual protein is a crucial determinant of its rate of flow transport, in addition to its molecular weight (16). Impinging flow from a micropipette positioned close to a supported lipid bilayer can concentrate or disperse lipid-anchored proteins in a similar way (17). If very high shear stress is applied to supported lipid bilayers, inter-leaflet slip between the upper and lower lipid leaflet occurs in addition to protein transport across the surface (18, 19).

Electric fields can also move proteins laterally across membranes. Charged proteins and lipids can be driven by an electric field parallel with the membrane in model glass-supported lipid bilayers (20–22) and in bilayer patches formed from plasma membrane vesicles (23, 24). Similar lateral transport of charged membrane components occurs on the surface of living cells. A variety of different cell types display galvanotaxis, or migration aligned with electric fields. Electrophoresis of charged surface proteins has been proposed to direct this movement (25, 26). Recently, the Theriot group identified a novel electric field sensor protein called Galvanin, a strongly negatively charged transmembrane protein responsible for electric field-directed migration of keratocytes (27, 28). Within a few minutes after a field is applied, this protein accumulates at the positive end of the cell and the dynamics of both accumulation and subsequent changes in the direction of the cell’s motion are consistent with protein diffusion rates (29). Heparan sulfate is a polysaccharid that is a major component of the glyocalyx and is attached to many cell surface proteoglycans. Electrophoresis of heparan sulfate on multiple cell types is correlated with electric-field aligned migration (30).

Flow-mediated transport of surface proteins on living cells has not been observed so often. We are aware of only one clear-cut example: the unicellular hemoflagellate *Trypanosoma brucei* generates small flows as it swims through the bloodstream of its host. The resulting drag force on host immunoglobulins bound to lipid-anchored surface glycoproteins moves them to the posterior end, where they are endocytosed (31).

Endothelial cell surface proteins are sometimes observed to relocate in the downstream direction, consistent with flow transport. In 2013, Zeng et al. used immunolabeling to identify flow-dependent redistribution of heparan sulfate on a confluent culture of rat fat pad endothelial cells. Cells that were subjected to 10 minutes of shear stress displayed concentration gradients of heparan sulfate (32); at longer times, heparan sulfate was found primarily at intercellular junctions. Similar gradients formed on bovine endothelial cells after flow was applied for 30 minutes (33). This finding was specific to heparan sulfate; surface distributions of other glycocalyx components did not change under flow. The authors concluded that the observed gradients resulted from movement of a sub-population of a particular proteoglycan, glypican-1, along with its bound heparan sulfate.

Transmembrane proteins also appear to move downstream under flow. Vascular endothelial protein tyrosine phosphatase (VE-PTP) is a single-pass transmembrane protein expressed in endothelial cells. Mantilidewi et al. used immunofluorescence to observe shear-stress induced redistribution of VE-PTP in mouse brain endothelial cells, human umbilical vein endothelial cells, and human embryonic kidney cells (34). A subsequent study by Shirakura et al. identified similar polarized distribution of VE-PTP in endothelial cells obtained from whole mounts of mouse aorta (35). The gradients appear similar to those that form on nonliving model systems under shear stress. However, these observations were both made using immunolabeling of fixed cells. With only static snapshots of the protein redistribution, it is difficult to distinguish active from passive transport.

Here, we imaged rapid, reversible redistribution of two different lipid-anchored proteins on the surface of living cells under applied shear stress (figure 1). We expressed glycosylphosphatidylinositol-green fluorescent protein (GPI-GFP) and a fluorescent construct of glypican-1 and GFP (glypican-1-GFP) (figure 1 C) at the surface of COS-7 cells. We chose glypican-1 because it has been proposed as a possible endothelial cell flow sensor in previous work (36). We observed that cell-wide, lateral concentration gradients formed within minutes after flow was started, remained constant while flow was on, and disappeared after flow was stopped. Steady-state concentration profiles were well fit by exponential functions, as expected for passive transport of diffusing membrane proteins. Using our previously developed image analysis methods, we determined the hydrodynamic force on each protein (16). When we labeled each protein with an antibody to GFP, we observed a significant increase in the hydrodynamic force and in the resulting flow transport. We did not observe asymmetric actin reorganization on the same time scale as the GPI-anchored protein transport. Our observations are consistent with passive flow transport of GPI-anchored proteins on living cells.

**Fig. 1.**
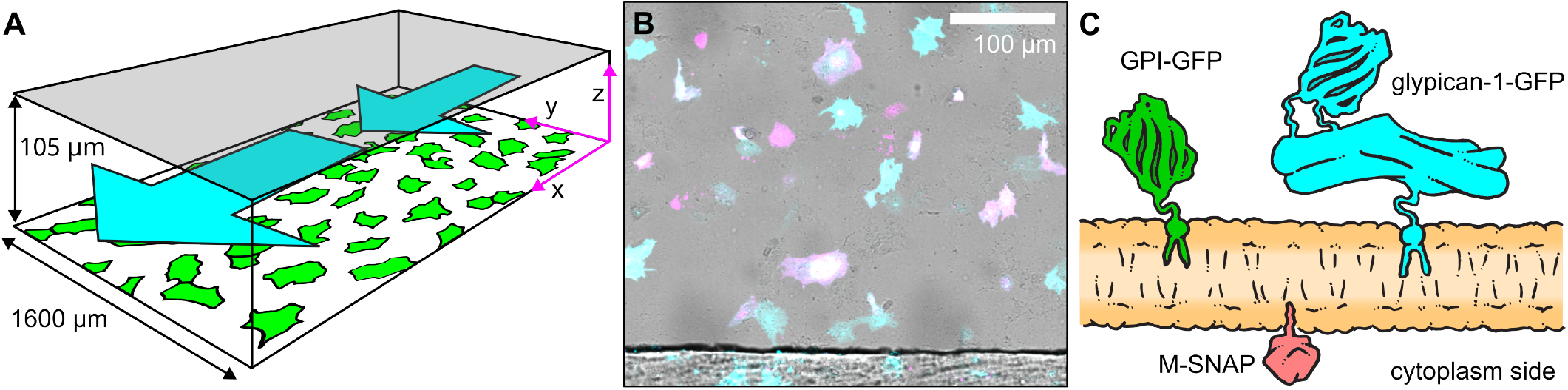
Experimental overview. A) COS-7 cells were seeded on a coverslip, then assembled into a rectangular flow chamber with the coverslip forming the lower surface. Flow was applied in the *x* direction using a syringe pump. B) Composite image showing COS-7 cells expressing GPI-GFP (cyan) and LifeAct (magenta) after assembly into a flow channel imaged in brightfield (grayscale). One edge of the silicone gasket is visible at the bottom of the image. C) Illustration of the membrane proteins used in this experiment.

### Experimental Methods

#### Mammalian Cell Culture

We chose COS-7 cells for their flat morphology, good adherence under fluid flow, and ease of transfection. COS-7 cells were cultured in Cytiva HyClone Dulbecco Modified Eagle Medium (DMEM) with high glucose with L-glutamine and sodium pyruvate and supplemented with 10% fetal bovine serum (FBS), 100 units/mL penicillin and 0.1 mg / mL streptomycin. Media used for transfection did not include additional supplementation with FBS, penicillin, or streptomycin (referred to as no-additive media). The cells were cultured in a humidified atmosphere with 5% CO_2_ at 37 °C.

#### Glypican-1-GFP construct

In order to obtain fluorescent glypican-1, we designed a construct called glypican-1-GFP. To preserve protein processing and trafficking, mid-sequence insertion of the fluorescent label was required in order to avoid alteration of both the N-terminal signal peptide and the C-terminal GPI anchoring site of glypican-1. We chose the disordered region between residues 339-364 since it is located opposite to the GPI anchor site (see supplementary figure 1). Insertion of the eGFP sequence between residues 355 and 356 was successful. Since we did not want the eGFP to constitute the C-terminus, the stop codon of the eGFP was removed. The optimized glypican-1-GFP sequence was produced and cloned into a pcDNA3.1(+) vector by GenScript Inc. (Piscataway, NJ, USA).

#### Transfection

40 mm round glass coverslips were placed in 60 mm culture dishes and treated with Cultrex Poly-L-Lysine (R & D Systems, Minneapolis, MN) for 2 h at 37 °C, then washed with 1X PBS before seeding COS-7 cells in the culture media. Cells were grown to approximately 90% confluency prior to transfection. At least one hour before transfection reagents were added, media was removed and cells were washed with PBS before replacing 4 mL no-additive media over cells. Transfections were done using Lipofectamine 2000 Transfection Reagent (Invitrogen) and following manufacturer recommendations. As recommended for 60 mm dishes, 20 *µ*L lipofectamine was diluted to 500 *µ*L with no-additive media. For co-transfection, 8 *µ*g DNA #1 and 8 *µ*g DNA #2 were diluted to 500 *µ*L with no-additive media. After 5 minutes at room temperature, tubes of diluted lipofectamine and DNA were mixed and incubated together for 30 minutes at room temperature. The 1 mL mixture was added dropwise over cells to transfect and gently rocked to mix before incubating overnight at 37 °C with 5% CO_2_. For co-transfections, DNA #1 was either glypican-1-GFP pcDNA3.1+ or pCAG:GPI-GFP. pCAG:GPI-GFP was a gift from Anna-Katerina Hadjantonakis (Addgene plasmid # 32601 (37)). DNA #2 for co-transfections was either M-SNAP, a gift from Andrea Stoddard and Sarah Veatch, or pLifeAct_mScarlet-i_N1, a gift from Dorus Gadella (Addgene plasmid # 85056 (38). The SNAP-tag sequence (39) was appended to the sequence for M, a short N-terminal region of the Src15 gene that contains a myristoylation site and a short polybasic region (40). Transfected COS-7 cells were used for flow experiments the following day. Cells appear sparse in our fluorescence images for two reasons: first, transfection efficiency was generally low, typically between 10 and 20%, and second, some cells detached from the surface between transfection and imaging.

#### Imaging

Flow buffer (10 mM HEPES, 135 mM NaCl, 5 mM KCl, 10 mM glucose, 1 mM MgCl, 1.5 mM CaCl_2_, 1% fetal bovine serum (FBS) and 0.5% bovine serum albumin (BSA) at pH 7) was prepared on the day of imaging. We used a membrane-permeable SNAP-Cell TMR-Star to label the M-SNAP protein at the inner plasma membrane of COS-7 cells. We incubated cells in a labeling solution of TMR-Star at 1*µ*M in DMEM with 10% FBS at 37 °C and 5% CO_2_ for 30 minutes immediately before imaging. For antibody labeling, GFP Polyclonal Antibody pre-labeled with Alexa Fluor 594 (Thermo Fisher Scientific, Waltham, MA) was diluted in flow buffer and cells were incubated for 10 minutes just prior to imaging. To minimize bubbles, flow buffer lacking BSA and FBS was used to wash the cells before assembling the coverglass into the flow chamber for imaging.

Confocal imaging and flow experiment: The coverglass with cells was carefully assembled in a flow chamber (Bioptechs, Butler, PA) composed of a silicone flow gasket with nominal height 100 *µ*m and width 1500 *µ*m separating the coverslip from an upper glass slide. The flow chamber was maintained at 37 °C. We used a syringe pump to inject flow buffer at the desired flow rate. We imaged the cells using a spinning disk confocal fluorescence microscope with a computer-controlled motorized stage (Intelligent Imaging Innovations, Denver, CO). Expression of both glypican-1-GFP and GPI-GFP was highly variable, so we selected cells with good signal and performed a timelapse acquisition at multiple positions. At each location, we collected confocal stacks using a 63X oil objective and excitation with 488 nm and 561 nm lasers sequentially. Movies were recorded for 40-60 minutes, with 2-4 minute intervals between time points.

#### Image analysis

##### Cell selection and region of interest

COS-7 cells displayed a mostly flat morphology with cell area ranging between approximately 600 and 1800 *µ*m^2^. Except in the region close to the nucleus, the apical and basal membranes were so close together that they could not be distinguished (figure 2). Cross-sections of the confocal stack show that most of the fluorescence from GPI-GFP, glypican-1-GFP, and the M-Snap-tag was located at the membrane; however, we also frequently observed bright fluorescence near the nucleus that was located in the cytoplasm (Figure 2 A-C). This perinuclear region most likely includes the Golgi, typically localized in the upstream direction in endothelial cells subjected to blood flow (41). In order to limit our analysis to the external, membrane located proteins, we manually selected regions of the cell that appeared flat, close to the coverslip, and free from high-intensity internal fluorescence (figure 3 A-C). Cell edges moved during the acquisition time (see supplementary movies 1 and 2), so we also limited our analysis to cells which did not show extensive, directional movement and chose regions that remained inside the cell boundaries throughout the acquisition.

**Fig. 2.**
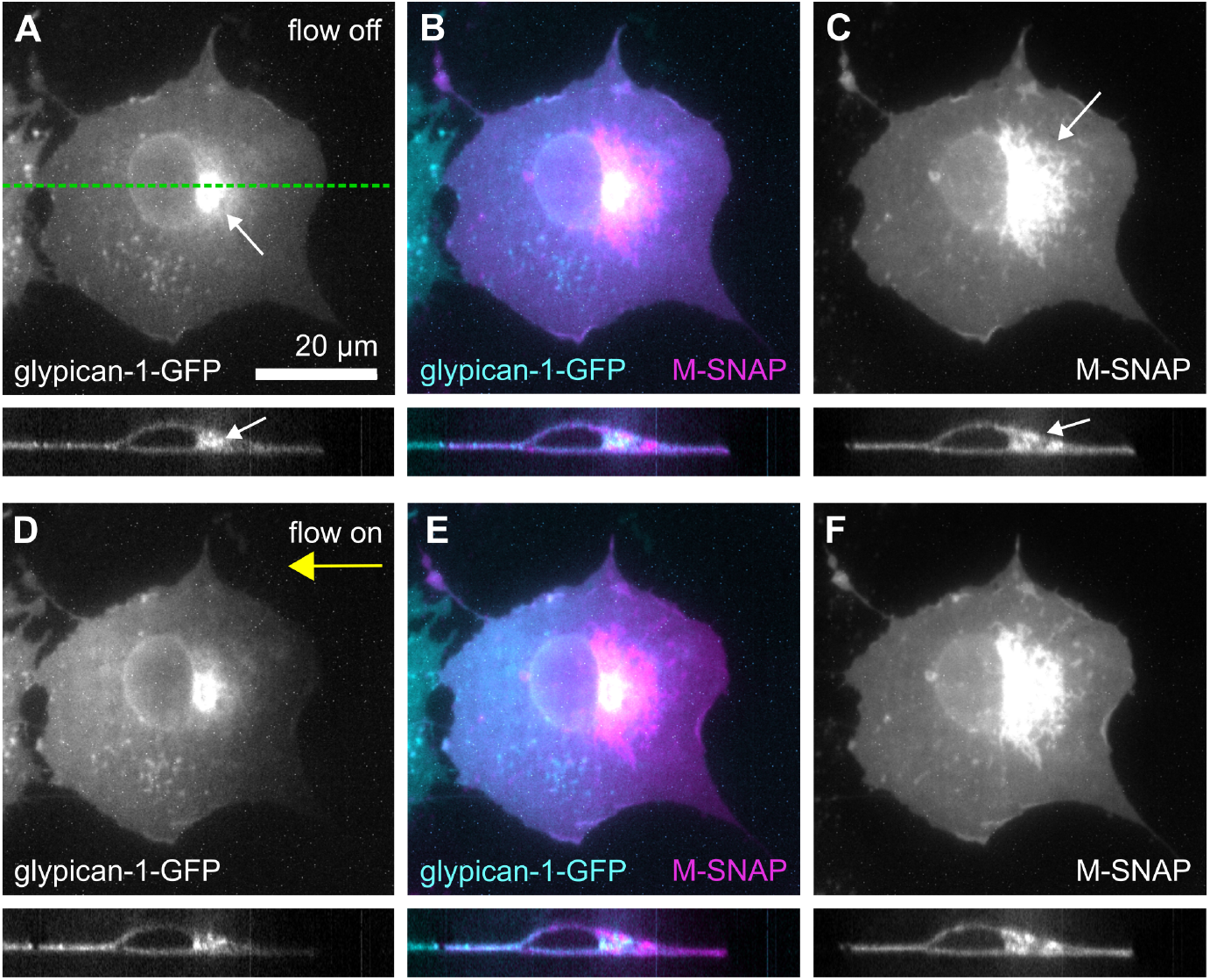
Evidence for flow transport of GPI-anchored proteins. COS-7 cells were doubly transfected with glypican-1-GFP (A) and M-SNAP (C), then labeled with cell-permeable label SNAP-Cell TMR-Star. Panels A-C and D-F show sum projections of confocal stacks of a typical cell. Below each projection is the *xz* plane cross section located at the green dotted line in panel A. The cross sections show that both fluorescence signals are located at the plasma membrane, with the exception of bright perinuclear regions (small arrows). In static conditions, both proteins were distributed evenly across the surface of the cell (A-C). After 3.3 Pa of shear stress was applied for 4.3 minutes (panels D-F, the membrane-located glypican-1-GFP, but not the M-SNAP, shifted in the direction of flow (large arrow). Internal fluorescence did not change position under flow.

**Fig. 3.**
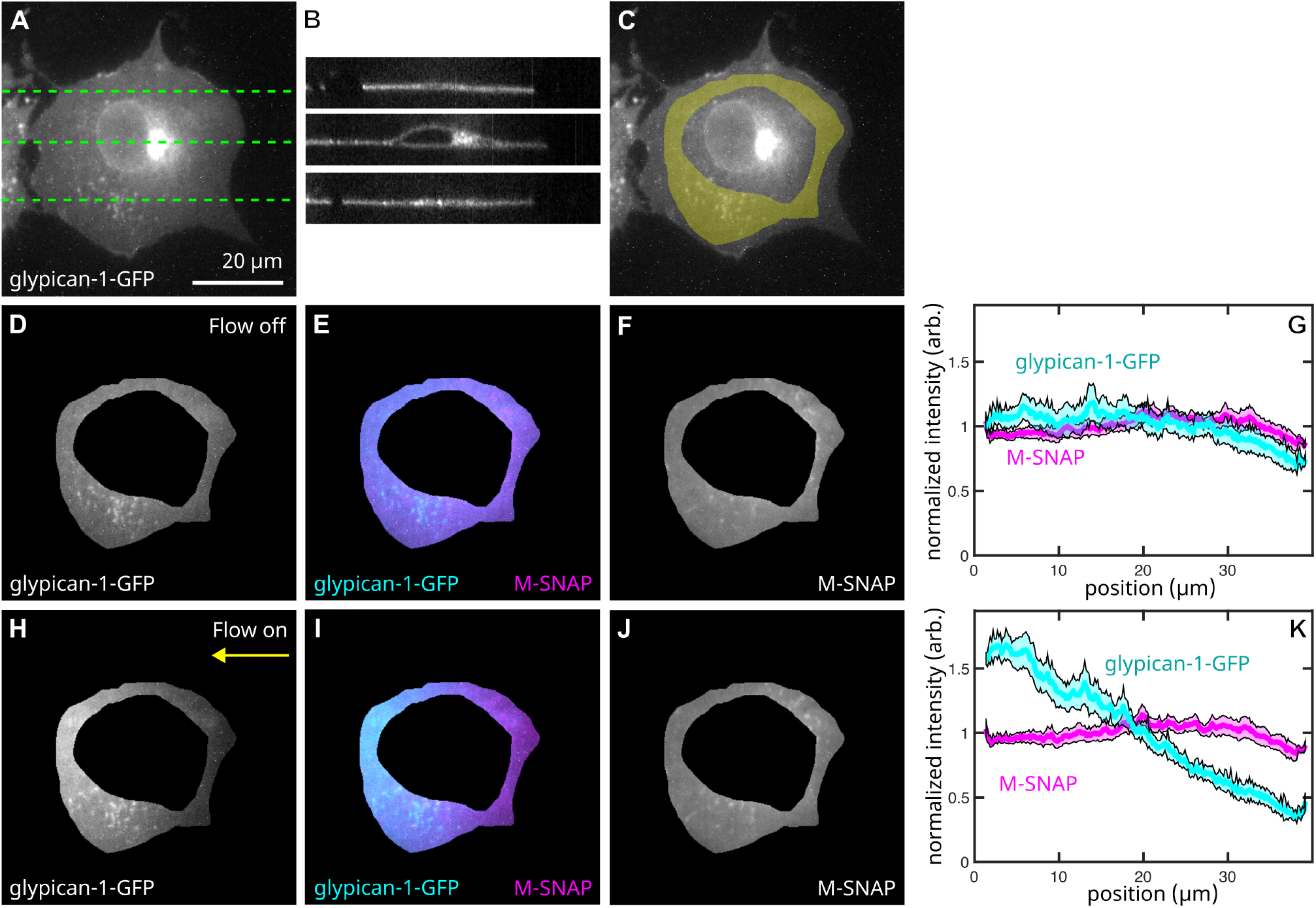
Isolation of membrane-located fluorescence. We hand-selected regions located away from the nucleus and that were inside the cell boundaries at all time points. The sum projection of a confocal stack (A) with *xz* cross sections of the confocal stack located at the dotted lines (B) show that fluorescence from the upper and lower membranes could be distinguished only near the nucleus. An example selected region is overlaid with the cell image in C. Fluorescence from glypican-1-GFP (D, E) and from M-SNAP (E, F) in the selected region was homogenous in the absence of flow. Panels H-J show the fluorescence in the same region after 3.3 Pa of shear stress was applied for 4.3 minutes. We vertically averaged fluorescence in the selected region and the average is plotted vs. position in panels G and K without and with flow, respectively. Shaded areas are one standard deviation wide. For ease of comparison, intensity profiles in G and K were separately normalized.

Fluorescence from the chosen region in each cell was collected by making a sum projection of the confocal stack. Since we collected the stacks at the optimal Nyquist spacing in the z-direction, we expect that the sum projection intensity is proportional to the total concentration of fluorescent protein in each pixel.

##### Normalization

unlike the supported lipid membranes that we have analyzed previously, cell membranes are dynamic and unlikely to be flat on the nanometer scale (42). Three-dimensional membrane topography can interfere with fluorescence-based determination of membrane protein concentration (43). To correct for this, we monitored a third membrane protein, M-SNAP. Since this protein is located on the cytoplasmic side of the membrane, we expect that it will not be transported by flow; we use it to monitor changes in membrane density from both large-scale changes in cell shape and sub-micron membrane wrinkles or invaginations. We divided the GFP fluorescence signal by the M-SNAP signal at each pixel and time point to obtain GFP concentration relative to the amount of membrane for our analysis (figure 4). We were suprised to find that although this normalization impacted the results for individual experiments, average results were indistinguishable with and without normalization (Supplementary figure 2). This suggests that in the flat regions we chose for analysis, sub-micron membrane curvature was negligible and did not change significantly under shear stress. We used normalized intensity to determine the hydrodynamic force and diffusion coefficient for both glypican-1-GFP and GPI-GFP (data shown in figure 6 and Table 1). In subsequent experiments including either LifeAct or antibody, we did not include M-SNAP transfection, labeling, or normalization.

**Table 1.** Summary of results

| Protein | Diffusion coefficient ($\mu\text{m}^2/\text{s}$) | HF (fN/Pa) | Hydro. Area ($\text{nm}^2$) | Molecular Weight (kDa) |
| --- | --- | --- | --- | --- |
| GPI-GFP | $0.88 \pm 0.54$ | $0.020 \pm 0.009$ | $20 \pm 9.4$ | 27 |
| GPI-GFP + antibody | $0.33 \pm 0.1$ | $0.050 \pm 0.042$ | $50 \pm 42$ | 177<sup>a</sup> |
| glypican-1-GFP | $1.04 \pm 0.64$ | $0.105 \pm 0.027$ | $105 \pm 27$ | 89<sup>b</sup> 105<sup>c</sup> |
| glypican-1-GFP + antibody | $0.21 \pm 0.2$ | $0.189 \pm 0.143$ | $189 \pm 143$ | 240<sup>a,b</sup> 255<sup>a,c</sup> |

<sup>a</sup> Assuming an approximate antibody molecular weight of 150 kDa.
<sup>b</sup> As predicted from sequence, neglecting any glycosylation.
<sup>c</sup> As measured (see supplementary material).

**Fig. 4.**
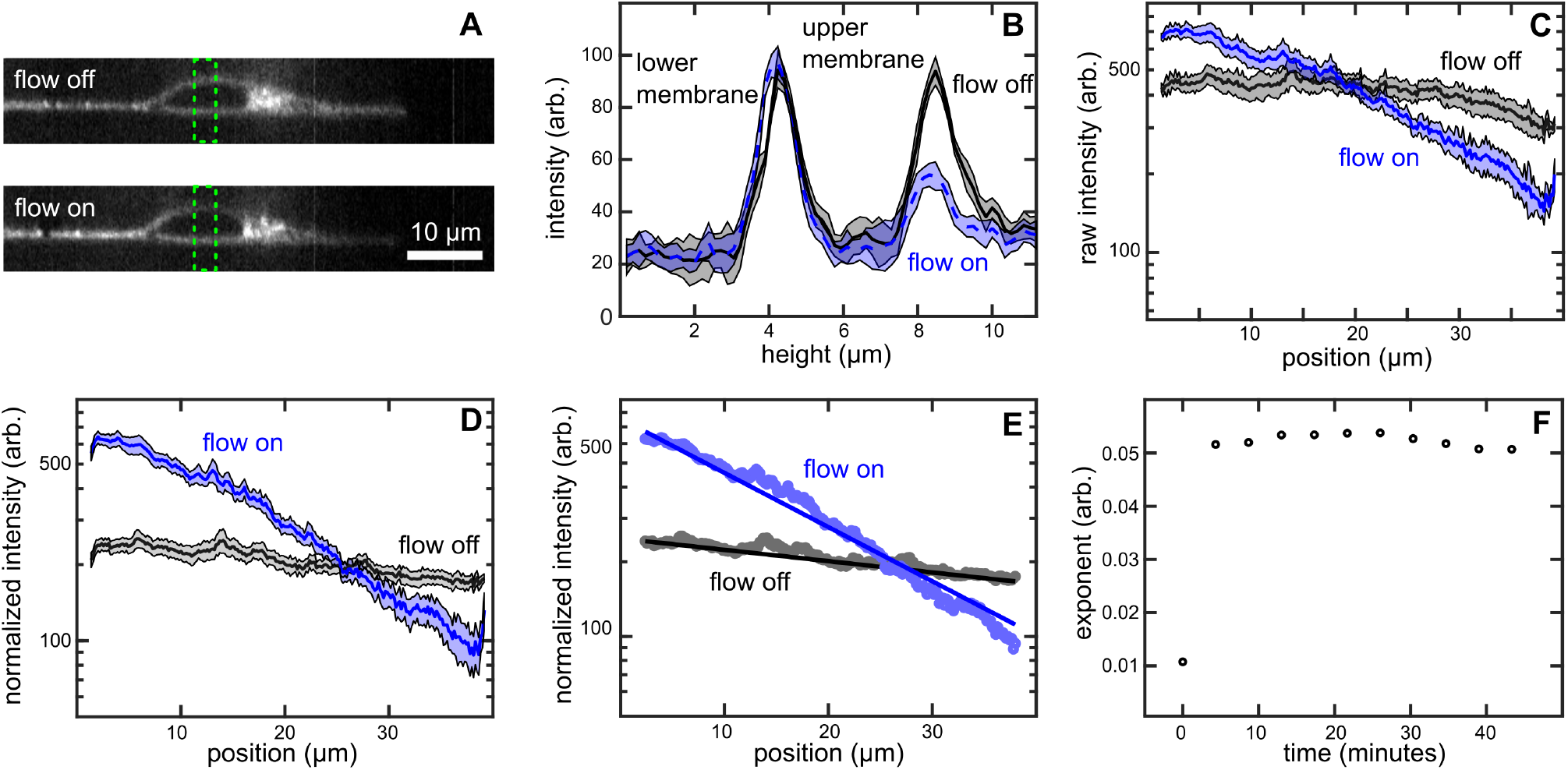
Isolation and normalization of signal from the apical membrane. Panel (A) shows an *xz* cross-section through the same cell shown in figure 3 before and after flow was applied. We plotted the average intensity inside the green dotted boxes shown in panel A for the same cell before flow was applied (solid line) and after 3.3 Pa of shear stress was applied for 4.3 minutes (dashed line). Under flow, the upper (apical) membrane signal decreased while the lower (basal) membrane signal remained approximately constant (B). Panel (C) shows the raw fluorescence intensity profile of the sum projection in the flat region (identical to those illustrated in figure 3). In order to isolate signal from the upper membrane, we subtracted half of the intensity of the first frame (recorded in the absence of flow) from each subsequent frame. We then divided the GFP signal intensity by the M-SNAP signal (panel D). This subtracted and normalized profile was then fit to an exponential function as previously described (E). The resulting exponent remained constant over tens of minutes while flow was on (F). In panels B-D, the shaded areas are one standard deviation wide.

##### Removing intensity from the lower membrane

only proteins in the apical membrane are subjected to shear stress during flow, so we want to monitor signal from the apical membrane only. In regions where the upper and lower membrane were far apart, we observed that fluorescence from the lower membrane did not change under flow (Figure 4 A-B). We therefore assumed that intensity from the lower membrane stayed constant during our movies, and changes were due only to protein distribution in the upper membrane. We started each movie by imaging the cell under static conditions, so that proteins were distributed randomly over the cell. We assumed that half of the fluorescence intensity recorded in the first frame of each flow movie originated in the lower membrane. We therefore subtracted half of the intensity recorded in the first frame of the movie from all the subsequent frames. The remaining intensity was then averaged in the *y* dimension to obtain the intensity profile *i*(*x*) (figure 4 D) and this profile was analyzed over time as previously described (figure 4 E) (16) to determine the hydrodynamic force applied by flow to the proteins. Briefly, we fit the intensity profile to an exponential function in which the exponent represents the ratio of flow-driven drift velocity *v* to diffusion coefficient *D*:

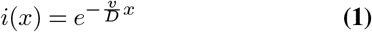

(the negative sign on the exponent indicates flow is in the -x direction, right to left in our images). We assume that under flow a protein reaches a terminal velocity 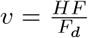 where *HF* is the hydrodynamic force on the protein, and *F_d_* is the sum of all membrane and fluid drag forces. Since the diffusion coefficient is the ratio of *k_B_T* to *F_d_*, we obtain the hydrodynamic force directly from the exponential fit coefficients by multiplying 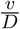 by *k_B_T* .

##### Diffusion coefficient measurement

after flow was stopped, we observed relaxation of flow-induced concentration gradients (figure 5 A-C). The time constant of this relaxation is related to the protein diffusion coefficient D by the expression (15, 44):

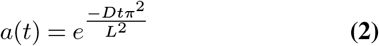

where *a* is the amplitude of the first cosine mode of the intensity profile and *L* is the diameter of the analysis region in the flow direction. Data were analyzed as described previously with the modification that intensity profiles were fit to a cosine function to obtain time-dependent amplitudes (figure 5 D, E). This allowed us to complete the fit even in cases where the selected region did not span the entire cell. The accuracy of our diffusion measurement was limited in these experiments by the slow frame rate required to collect stacks at multiple locations inside the flow channel; since GPI-anchored protein diffusion coefficients are high, gradients disappeared after only a few frames.

**Fig. 5.**
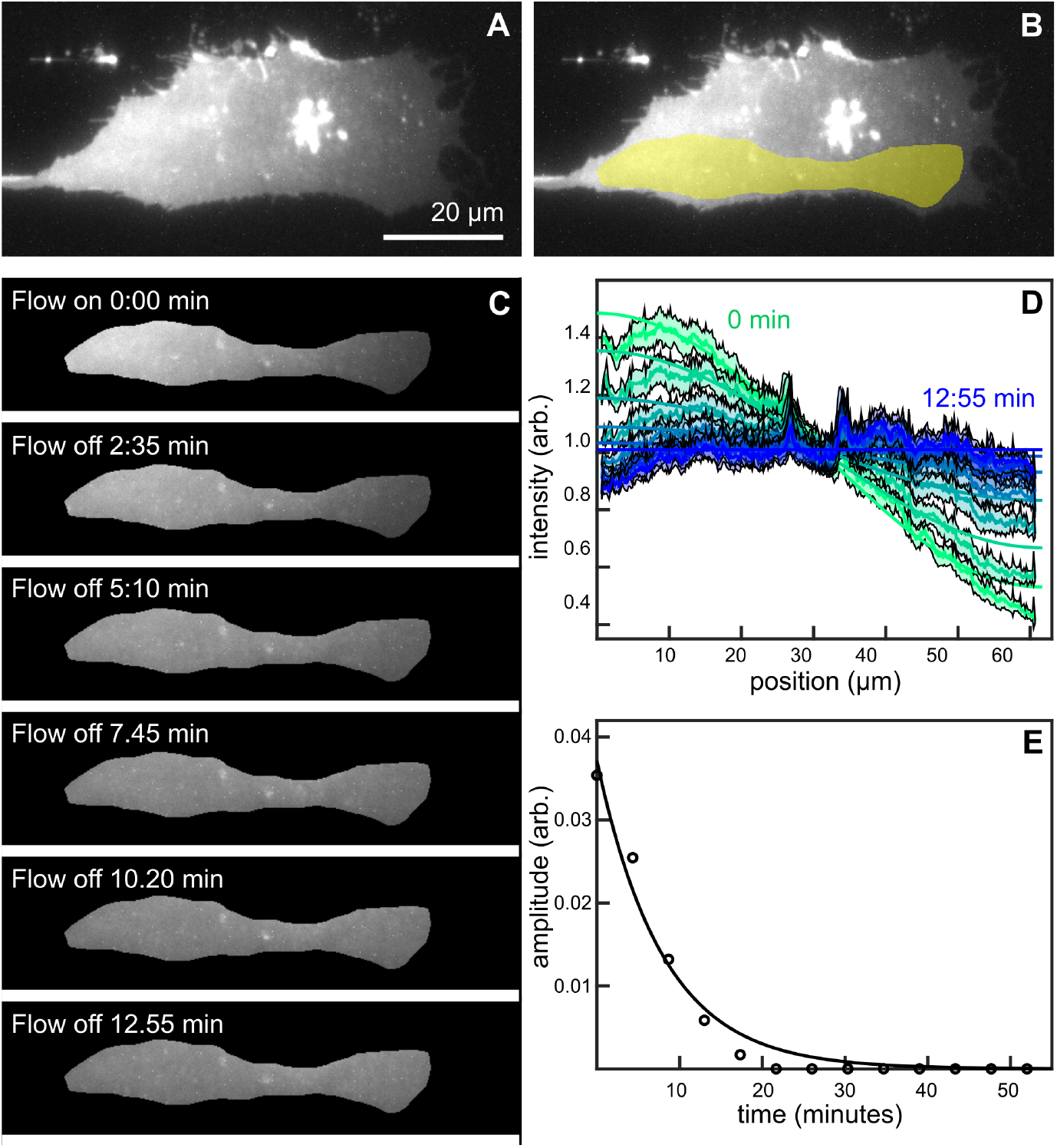
We measured diffusion by observing gradient relaxation dynamics after flow was stopped. Panel (A) shows GPI-GFP fluorescence on a COS-7 cell under 6.65 Pa of shear stress. We selected a flat region of the cell (B) and observed disappearance of the intensity gradient after flow was turned off (C). We generated the intensity profile from the selected region, normalized total fluorescence and fit the intensity profiles to cosine functions (D). Shaded regions are one standard deviation wide. The diffusion coefficient can be determined by fitting the resulting amplitude time decay.

##### Shear stress calculation

the silicone gasket we used with our flow chamber forms a wide, flat rectangular channel with nominal height 100 *µ*m and width 1500 *µ*m. In such a channel, the surface shear stress at the lower coverslip varies by less than 4% in all regions more than 100 *µ*m from the channel edges. We chose cells located at least 150 *µ*m from an edge and calculated the average shear stress in this region using the channel dimensions, flow rate, and viscosity (16, 45). We prepared flow buffer, heated it to 37 °C in a water bath, and measured the viscosity using an Ubbelodhe viscometer. We determined channel height by repeatedly assembling the flow channel with air inside and focusing on the inner coverslip surfaces to determine the distance between them, resulting in an average assembled height of 105 *±* 14 *µ*m and an average assembled width of 1600 *±* 70 *µm*. The COS-7 cells were mostly flat, with the maximum height in the nuclear region approximately 5-6 microns above the rest of the cell. We determined the point spread function of the microscope in the z-direction by scanning through a glass-supported lipid bilayer and used this to estimate that cell thickness in flat regions was 0.3 *µ*m or less. Since this thickness is much smaller than the uncertainty in channel height, we neglected it in our shear stress calculation.

## Results

We found that concentration gradient formation and dissipation were consistent with passive flow transport. In the absence of flow, all three proteins (GPI-GFP, glypican-1-GFP, and M-SNAP) were distributed evenly over the surface of each cell. Within minutes after flow onset, we observed downstream movement of both extracellular GPI-anchored proteins, but not of the cytoplasm-side M-SNAP protein. Both GPI-anchored proteins quickly formed steady state concentration gradients that were well fit by exponential functions, and the gradients stayed constant as long as the flow was on (figure (4 F, up to 50 minutes). For passive flow transport, we expect measured hydrodynamic force on the proteins to be linearly proportional to the applied shear stress, and this is what we observed (figure 6 A). Linear fits to the experimental data yielded the average hydrodynamic force. Although there was variation between individual cells, we found that the average hydrodynamic force on glypican-1-GFP was significantly larger than that on GPI-GFP (figure 6 A). When the flow stopped, gradients disappeared over 4-16 minutes due to diffusion. We determined average diffusion coefficients of approximately 0.88 *±* 0.54 *µm*^2^*/s* for GPI-GFP and 1.04 *±* 0.64 *µm*^2^*/s* for glypican-1-GFP (figure 6 B).

**Fig. 6.**
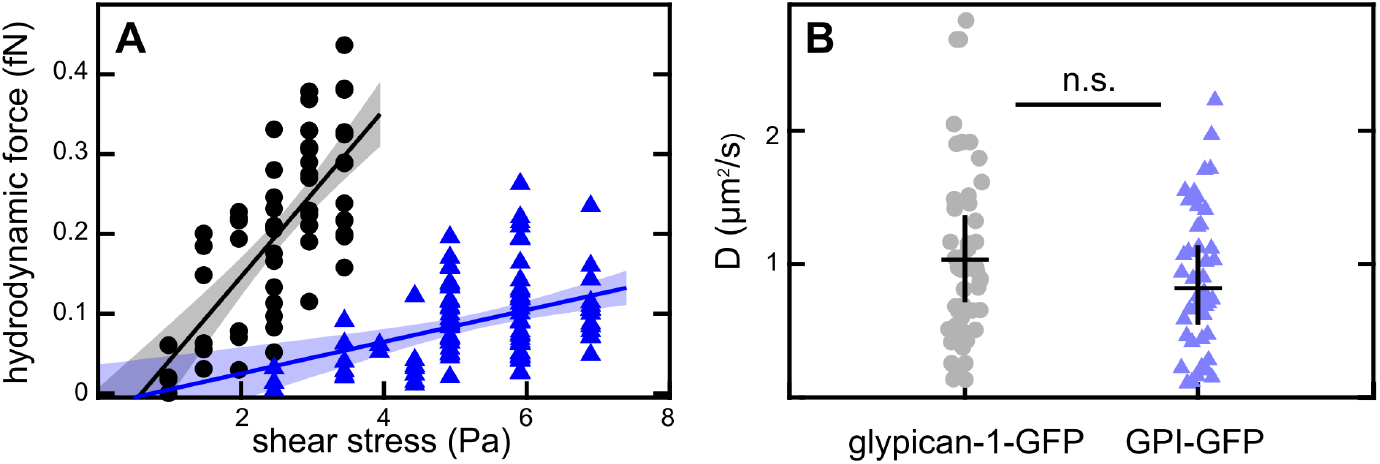
Collected results from many cells. A: Hydrodynamic force measurements recorded on 37 different cells expressing glypican-1-GFP (black circles) and 35 different cells expressing GPI-GFP (blue triangles). We fit each data set to a line; shaded regions indicate the 95% prediction intervals for the fit. The slope of the line yields the average hydrodynamic force on an individual protein per Pa of applied shear stress. Panel B: diffusion coefficients for glypican-1-GFP (gray circles) and GPI-GFP (blue triangles) were determined from gradient relaxation dynamics. Horizontal lines show the mean and vertical lines are one standard deviation in length.

If proteins are moving due to passive flow transport, increasing protein size should increase hydrodynamic force and the resulting transport under the same shear stress. We transfected cells with GPI-GFP and recorded videos of GFP transport under low shear stress, between 2.3 and 3.8 Pa, which we had previously found to result in minimal or no concentration gradient formation (figure 7 A, E, and I). We then incubated the cells with a fluorescent antibody against GFP and repeated the experiment, using the same flow rate. After antibody labeling, we observed small gradients in GFP fluorescence and large ones in the antibody signal (figure 7 B-K, see also supplementary movie 3). Observing the same cells before and after antibody labeling allowed us to be certain that variation between cells could not explain the difference in measured hydrodynamic force. We saw a substantial increase in hydrodynamic force after antibody labeling, from 0.020 *±* 0.009 fN to 0.050 *±* 0.042 fN. In other words, antibody labeling reduced the shear stress required to observe flow transport of GPI-GFP (figure 8 B). Antibody binding converted it from a protein that was effectively immobile under 3 Pa of shear stress to one that was readily mobile. The mobility increase was specific to antibody-labeled proteins (compare figure 7 panels J and K). Total antibody concentration was much lower than GFP concentration (see supplemental material for details).

**Fig. 7.**
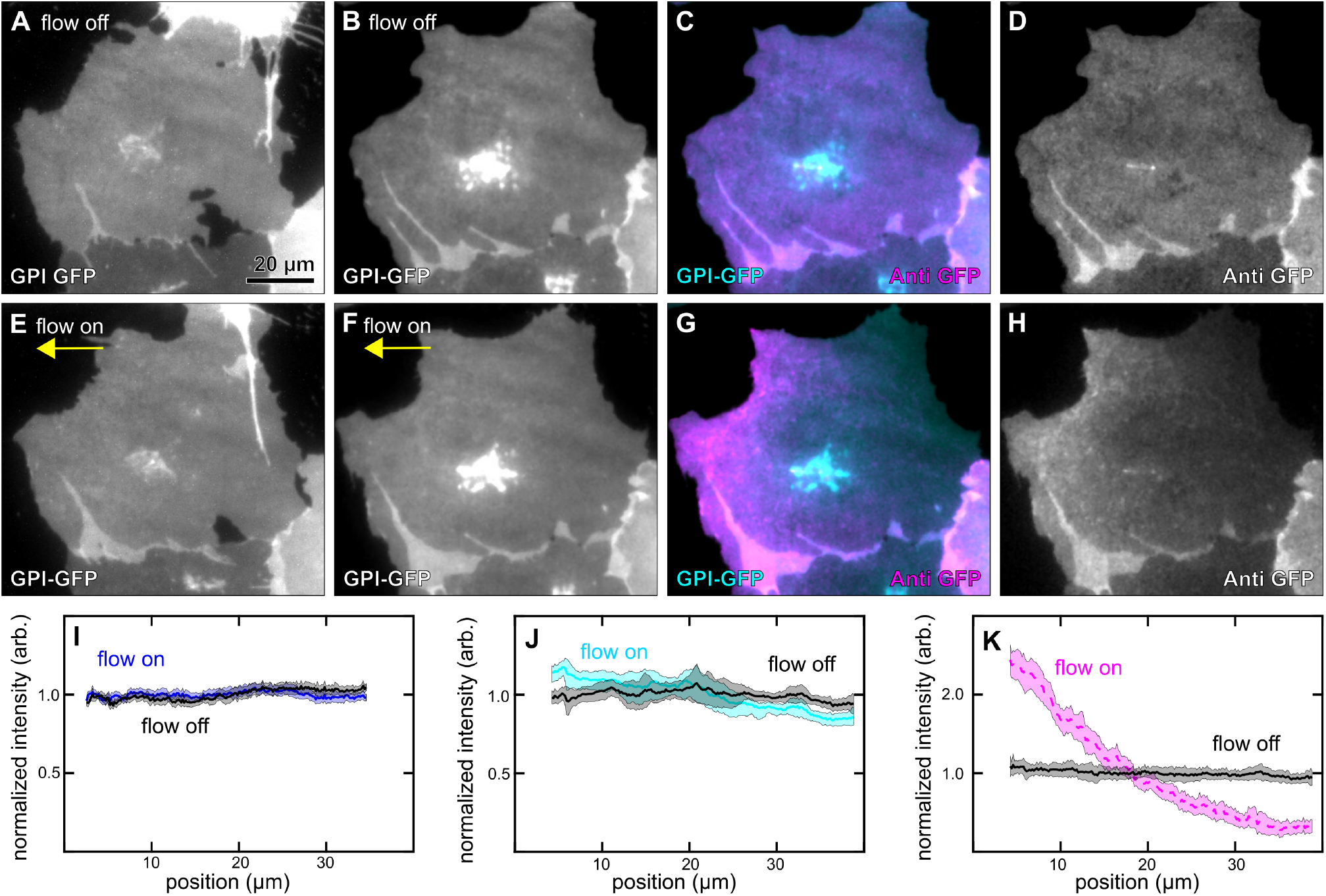
Antibody binding increased flow transport of GPI-GFP. We applied 2.37 Pa of shear stress to a COS-7 cell expressing GPI-GFP (panels A,E) and observed negligible change to the intensity profile (panel I). After incubating the same cell with an antibody to GFP, we applied the same shear stress for 15 minutes and observed a small gradient in the GFP intensity (B-C, F-G, and panel J) but a large gradient in the antibody intensity (C-D, G-H, and panel K). For ease of comparison, each profile was normalized separately before plotting. Shaded regions in I-K are one standard deviation wide.

**Fig. 8.**
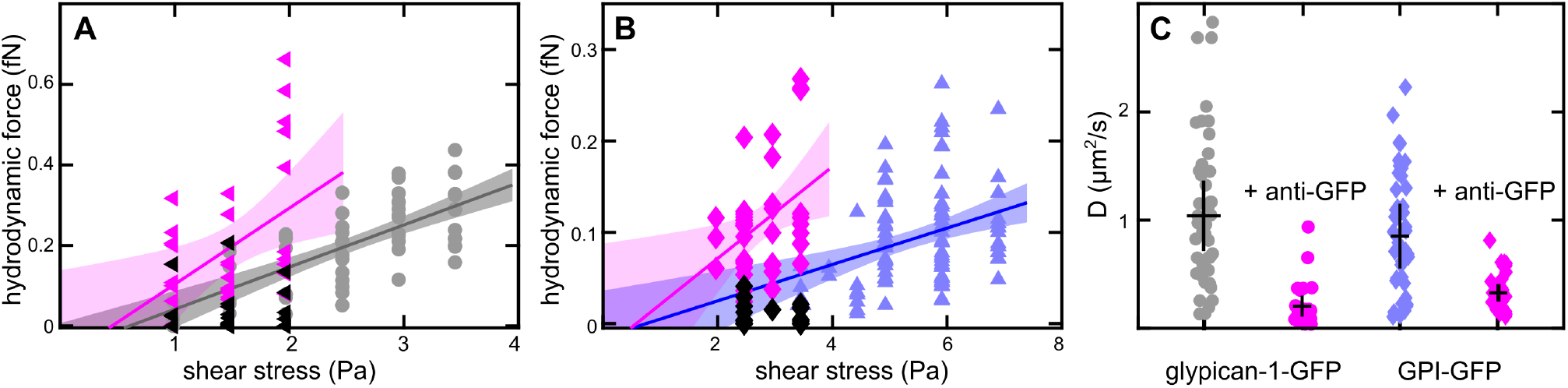
Antibody binding increased flow transport of both GPI-anchored proteins. In Panel A, the hydrodynamic force on 16 cells expressing glypican-1-GFP before (black triangles) and after (magenta triangles) antibody labeling is superimposed on the previous set of measurements (gray circles). Antibody fluorescence showed steeper gradients. Panel B: under low shear stress (2-4 Pa), flow transport of GPI-GFP was nearly undetectable (black diamonds), consistent with our previous determination (blue triangles). Antibody fluorescence on the same 37 cells showed strong gradients and average hydrodynamic force of 0.50 fN (magenta diamonds). Shaded areas in A-B indicate 95% prediction intervals for the linear fits. Panel C shows that the diffusion coefficients recorded for the fluorescent antibody complexes were smaller than those recorded for the proteins alone. Horizontal lines show the average value and vertical lines are one standard deviation in length.

We collected data before and after antibody labeling on multiple cells expressing either glypican-1-GFP (figure 8 A) or GPI-GFP (figure 8 B). In both cases, we observed an increase in hydrodynamic force (the increase was only statistically significant for the GPI-GFP-antibody complex). After stopping flow, we also observed a sharp reduction in the diffusion coefficients for the antibody complexes relative to the proteins in the absence of antibody (figure 8 C).

We co-transfected cells with GPI-GFP and LifeAct in order to simultaneously monitor GPI-GFP flow transport and actin filament location (figure 9, supplementary movie 4). We applied flow for approximately 16 minutes, and did not observe redistribution of actin in the flow direction.

**Fig. 9.**
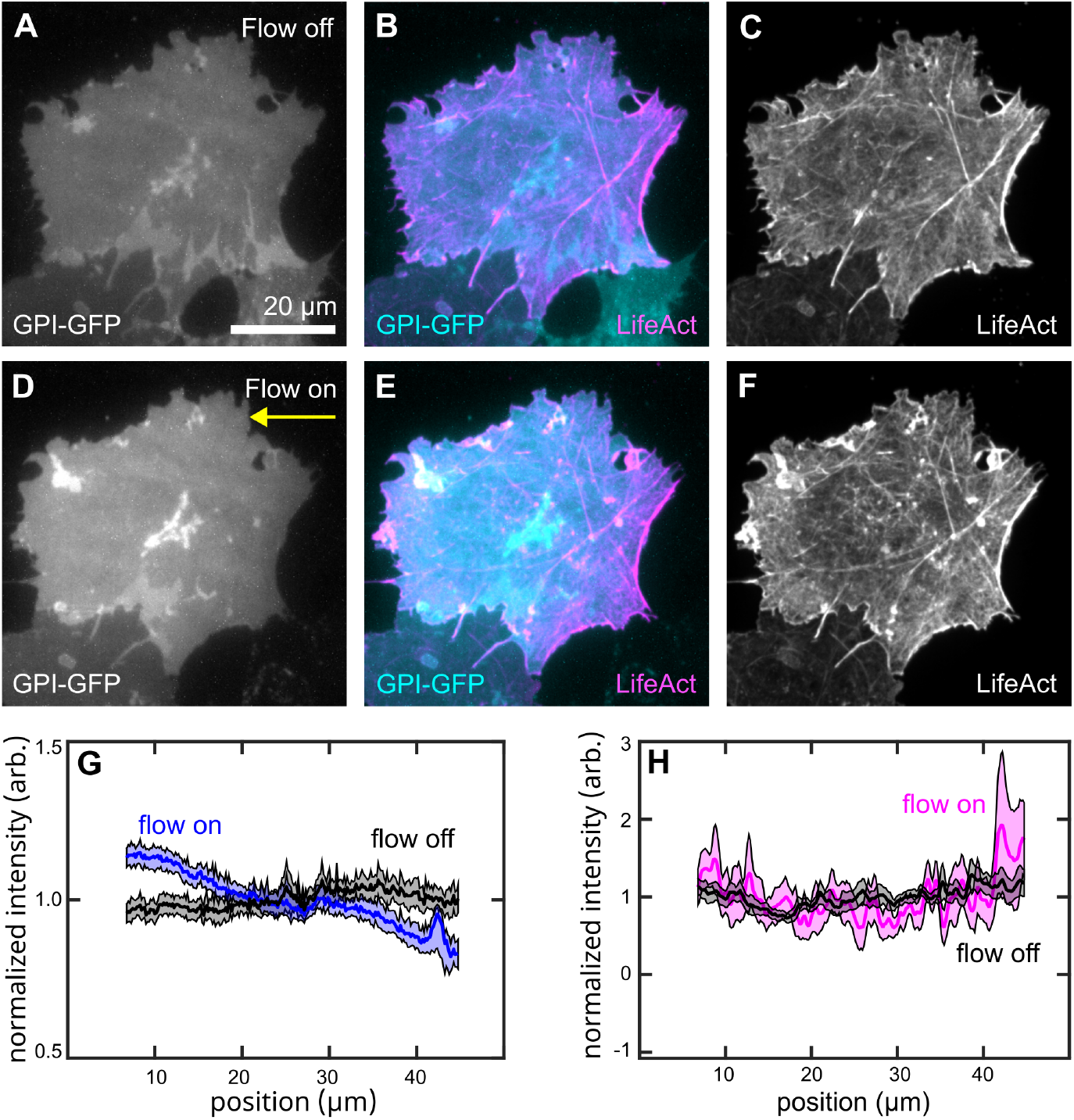
Actin distribution did not become asymmetric under flow. COS-7 cells co-transfected with GPI-GFP and LifeAct were subjected to 6.65 Pa of shear stress. GPI-GFP sum projections (A-B and D-E) show that GFP moved downstream after 9.5 minutes under flow. LifeAct maximum intensity projections (B-C and E-F) do not show motion in the downstream direction. This is visible in the average intensity profiles generated from the same region for GPI-GFP (panel G) and LifeAct (panel H). Shaded regions in G-H are one standard deviation wide.

## Discussion

Our results show that movement of GPI-anchored proteins on the surface of living COS-7 cells is consistent with passive flow transport. Specifically, we find that exponential steady-state concentration gradients form under flow, and that the dynamics of concentration gradients correspond to applied flow timing and magnitude. Hydrodynamic force and drift velocity are proportional to protein molecular weight and labeling proteins with an antibody results in increased hydrodynamic force and more efficient flow transport.

We can calculate the shear-induced drift velocity of our proteins by multiplying our measured exponential fit coefficient by the diffusion coefficient; using the averaged values we determined for each protein, we estimate that under one Pa of shear stress the drift velocity of GPI-GFP is 4.1 *±* 3.2 nm/s, and that of glypican-1-GFP is 25.5 *±* 17.3 nm/s. For both proteins, random diffusive motion over the course of a second is nearly an order of magnitude larger than motion in the flow direction. However, over longer times, flow-mediated drift becomes as important as diffusion. A protein traveling at 25.5 nm/s would take approximately 9 minutes to traverse the width of a 40 *µ*m cell, consistent with our observation that visible concentration gradients formed within a few minutes after flow was applied.

When we compared our results to our previous experiments on model membranes, measured hydrodynamic forces appear lower than expected. For example, we previously determined that one Pa of shear stress applied 0.054 *±* 0.018 fN to monomeric streptavidin, a 14.2 kDa protein, attached to a glass-supported lipid bilayer (16). In our current measurements of the motion of GPI-GFP on COS-7 cells, we found that one Pa of shear stress resulted in only 0.020 *±* 0.010 fN, despite the fact that GFP is larger (27 kDa). Similarly, the total hydrodynamic force on glypican-1-GFP in cells was similar to that found for tetrameric streptavidin on supported lipid bilayers, despite the fact that the molecular weight of glypican-1-GFP is significantly larger (89 kDa excluding glycosylation, compared with 52 kDa for streptavidin). The most likely explanation for these findings is that proteins on cells were partially shielded from the flow by the glycocalyx.

We used our results to make a rough estimate of the glycocalyx thickness. The hydrodynamic force on an object in shear flow is proportional to its hydrodynamic area, which depends both on its size and on its shape in the direction perpendicular to the membrane (46). In previous work, our measured hydrodynamic areas for lipid-anchored proteins correlated well to cylindrical shapes with reasonable molecular dimensions (16). GFP is a compact protein with a cylindrical structure, in which the radius is approximately one third of the height (47). We used the known molecular weight of GFP and an average value for protein density (48) to calculate its approximate volume. Using these two constraints, we model GFP as a cylinder with height 4.7 nm and radius 1.5 nm, perpendicular to the membrane surface. The predicted hydrodynamic area is 160 nm^2^ (46), much larger than our measured value of 20 *±* 10 nm^2^. That value is instead consistent with a cylinder with radius 1.5 nm and height 0.6 nm, as if the GFP molecule were embedded in a layer with an effective height of 4.1 nm (figure 10). Approximating the shape of glypican-1-GFP is more difficult, since all-atom simulations show that the shape of glypican-1 is irregular and that its orientation is highly dynamic due to the flexible link between the protein and its GPI anchor (49). However, we can make a conservative estimate for the hydrodynamic area of our glypican-1-GFP construct by approximating it as a sphere with radius 3.1 nm (using its measured molecular weight, and neglecting heparan sulfate chains). The hydrodynamic area of this sphere would be 312 nm^2^, higher than our measured value of 105 *±* 27 nm^2^. To model the effect of a 4.1 nm deep glycocalyx, we calculate the approximate hydrodynamic area of a hemispherical cap (50) with radius 3.1 nm and height 3.7 nm, giving a lower area of 82 nm^2^, closer to our measured value. Of course, this approximation oversimplifies by modeling the soft and variable glycocalyx as flat, impermeable surface. Still, it explains both the smaller observed force relative to model systems, and the trend in force with protein size (figure 10).

**Fig. 10.**
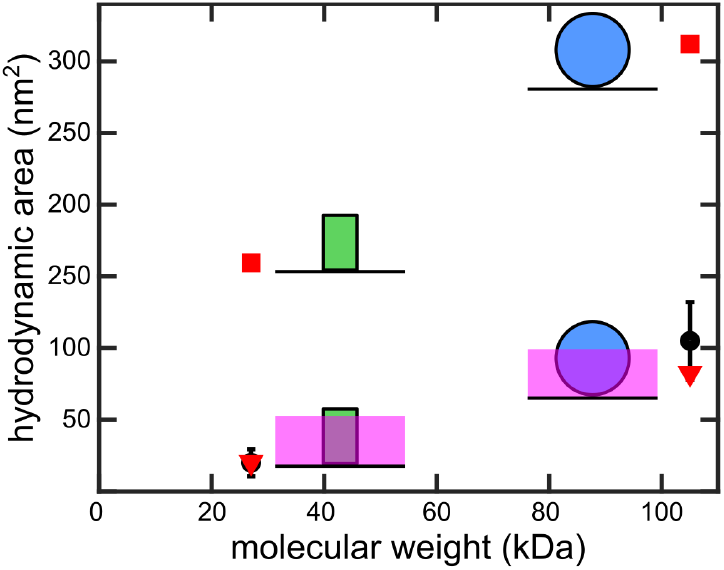
Comparison between measured and predicted hydrodynamic area. We plotted the average measured hydrodynamic area (black circles) for GPI-GFP and glypican-1-GFP versus the protein molecular weight (27 kDa for GFP, and 105 kDa for glypican-1-GFP). Vertical lines are two standard deviations in length. Using the molecular weight to predict total protein volume and approximating GPI-GFP as a cylinder (illustrated as a green rectangle) and glypican-1-GFP as a sphere (illustrated as a blue circle) located at the membrane surface, we predict much larger hydrodynamic areas for each (red squares). If we reduce the height of each shape by assuming it is partially embedded in a 4.1 nm thick glycocalyx (magenta), we predict hydrodynamic areas similar to our experimental measurements (red triangles).

Our estimate of glycocalyx thickness is much smaller than measured values of glycocalyx thickness on endothelial cells, which range from hundreds of nanometers to tens of microns (11). However, glycocalyx thickness varies in different cell types, and even in the same cells under different culture conditions. For example, endothelial cells grown on stiff substrates generate a thinner glycocalyx than those grown on soft ones (51, 52). We are not aware of any published measurements of COS-7 glycocalyx thickness, but it is likely to be thinner than that on endothelial cells. Optical profilometry and recent fluorescence measurements of molecular crowding provide evidence that surface proteins and glycans extend 3-20 nm from the membrane of multiple types of living cell (53, 54). Recent super-resolution imaging combined with fluorescent labeling of specific sugar residues shows glycocalyx components located 20-80 nm away from the membrane of breast cancer cells (55). Micropipette-based experiments yielded an estimate of 70 nm for the glycocalyx thickness of human mammary fibroblast cells (56). In addition to the glycocalyx, additional factors that could hinder flow-mediated protein transport include increased membrane drag on the GPI anchor arising from high membrane viscosity, crowding from mobile and immobile transmembrane proteins, or frictional drag from cytoskeletal protein contacts on the cytoplasmic side of the membrane. Additional experiments will be required to isolate these factors and to carefully model the flow shielding effects of membrane-attached polymers.

Estimating the effective size and shape of glypican-1-GFP is not trivial. Glypican-1 has three attachment sites for heparan sulfate. Heparan sulfate molecules can be large; estimates for the molecular weight of endothelial heparan sulfate range from 30 to 400 kDa (57, 58). Early studies in mouse embryo fibroblast cells found that the molecular weight of each glypican-1 heparan sulfate chain was greater than 60 kDa (59). While the heparan sulfate chains are attached so that they extend in opposing directions, they are also positioned close to the membrane, increasing the likelihood that other components of the glycocalyx shield them from shear stress. Furthermore, in live cells there is the possibility that some heparan sulfate chains interact with other proteins in the membrane, altering the drag forces on glypican-1-GFP. We ran a Western blot to determine the molecular weight of our construct when purified from COS-7 cells, and found that the apparent molecular weight was approximately 105 kDa, only slightly larger than the 89 kDa expected from the construct sequence (supplementary figure 5). This 105 kDa band probably represents the glypican-1-GFP construct carrying N-linked oligosaccharides but not full-length heparan sulfate chains (which may be sheared off of the protein during sample preparation).

The accuracy of our determination of diffusion coefficients was limited, since our acquisition frame rate was comparable to the time for a protein to diffuse across a cell. Despite this, our results fall within the range of previous determinations of diffusion coefficients for lipid-anchored proteins on living cell plasma membranes (between 0.25 and 1.75 *µm*^2^*/s*) (60– 63). We did not distinguish a significant difference between diffusion coefficients for GPI-GFP and glypican-1-GFP. For lipid-anchored proteins and nanoparticles, diffusion is dominated by the membrane viscosity and we expect the size of the aqueous domain to have only a moderate influence (64, 65). In addition, in membranes crowded with proteins, total protein concentration predicts diffusion coefficients better than protein size (66). Diffusion coefficients of antibody-labeled proteins were significantly smaller than expected (figure 8C), which may reflect cross-linking and aggregation (we observed that antibody fluorescence signal became more grainy over time, as visible in supplementary movie 4). Another possible explanation is that the antibody bound GFP in a configuration that resulted in it being very close to or partially embedded in the membrane. This would increase frictional drag and also lower the observed hydrodynamic force.

We made the assumption that shear stress at the surface of our cells was the same as that calculated for the lower surface of the microchannel. In reality, modeling shows that shear stress on rounded cells is directly proportional to the height of the upper membrane, with higher stress at the apex and lower stress in the valleys between cells (67). Our analysis was limited to the flat regions of the cells we identified, so the average shear stress value we used was most likely an overestimate. We did not control for each region’s position upstream or downstream from the cell nucleus, or for the presence or absence of surrounding cells. This topographical variation may explain some of the cell-to-cell differences in measured hydrodynamic force that we observed. Variation in the surface concentration of proteins could also impact our results. The concentration of both labeled and unlabeled surface molecules is expected to have a large impact on flow transport. Previous experiments showed that the effective hydrodynamic force on identically sized, mobile lipid-anchored proteins was highly dependent on their concentration because at high density, proteins partially shelter each other from the flow force (46). Variable expression of the GPI-anchored proteins could result in different apparent hydrodynamic forces.

Since all motor proteins are located in the cytoplasm, active transport of GPI-anchored proteins would have to be accomplished either via interactions with one or more transmembrane proteins, or via lipid coupling to transmit force from the inside to the outside of the membrane. The second mode is supported by experimental evidence that lipid interactions promote clustering of GPI-anchored proteins on the cell surface. Short, dynamic actin filaments generate nanometer-sized clusters and modify the dynamics of proteins via transmembrane coupling between inner leaflet phosphatidylserine and outer leaflet GPI anchors (68, 69). If actin filament-generated forces were responsible for the protein redistribution that we observe under flow, we would expect to observe asymmetric redistribution of actin, either before or at the same time as protein redistribution. To test this hypothesis, we simultaneously observed GPI-GFP and LifeAct under shear stress. A representative cell is shown in figure 9 and in supplementary movie 4. Under flow, actin intensity became more heterogeneous but did not develop asymmetry in the downstream direction.

As expected, we found that the larger protein glypican-1-GFP experienced a larger hydrodynamic force compared to the smaller protein GPI-GFP. Glypican-1-GFP concentration gradients were apparent on most cells under the shear stress typically found in blood vessels (1-2 Pa). To form similar gradients of GPI-GFP required significantly higher shear stress (5-7 Pa). An increase in downstream transport after antibody binding is a direct prediction from the model of passive flow-mediated transport. Conversely, a hypothetical active transport mechanism would require first detection of extracellular antibody binding, then signaling to achieve increased directed downstream transport. We observed that flow transport of the GFP antibody was larger than flow transport of the same proteins in the absence of antibodies (figure 8). In addition, the increase was specific to the small minority of GPI-anchored proteins with bound antibody (figure 7 B, F, J). For both proteins, the magnitude of the increase in hydrodynamic force was smaller than expected, given the large size of the GFP antibody. We believe that this is most likely explained by a tendency for the antibodies to crosslink multiple proteins and form aggregates over time. Both flow transport and diffusion would be strongly impacted by aggregation.

Although our experiments cannot completely rule out the possibility of directed active transport, passive transport by flow is the most straightforward explanation for our results. We speculate that observations of similar distinctive patterning of membrane proteins and oligosaccharides on flow-exposed cells (32–35) can also be explained in this way. VE-PTP has a sizeable extracellular domain consisting of 16-17 fibronectin-like domains, connected to the cytoplasmic domain by a single-pass transmembrane region (70). This structure is compatible with flow transport. Mantilidewi et al. determined that the cytoplasmic domain was not required for flow redistribution of VE-PTP, making it highly unlikely that motion was controlled or initiated via the interior of the cell. They further suggested that the large size of the extracellular domain explains why it is redistributed by shear stress, while other members of the same protein family are not (34). These findings are both consistent with direct flow transport of VE-PTP.

The primary protein components of the endothelial glycocalyx are syndecans and glypican-1, both carriers of heparan sulfate. Syndecans can also carry chondroitin sulfate (32). Unlike glypican, syndecans are transmembrane proteins. Ebong et al. showed that glypican-1, but not syndecan-1, is required for rapid flow activation of an enzyme, endothelial nitric oxide synthase, located in the cytoplasm. Both knockdown of glypican-1 and enzymatic removal of heparan sulfate prevented flow activation of endothelial nitric oxide synthase (36). Based on these results, the authors proposed that glypican-1 acts as a primary flow sensor in endothelial cells (36). Zeng et al. observed significant redistribution of heparan sulfate under flow and attributed it to movement of a sub-population of glypican-1; however, downstream movement of glypican-1 itself was not confirmed (32). This may reflect challenges specific to immunolabeling; glypican-1 is a difficult target for antibodies, possibly due to extensive glycosylation.

Parallel experiments recently completed by Yamashiro et al. describe observations of flow transport of multiple transmembrane proteins, showing that this mechanism of redistribution is not limited to lipid-anchored ones. In addition, their work shows variation in the flow mobility of one protein, GPI-GFP, when expressed in different cell types (71). We expect that if we observed flow transport of glypican-1 on endothelial cells, the magnitudes of hydrodynamic force and drift velocity would be smaller than the ones determined here due to the thicker endothelial glycocalyx.

COS-7 cells do not normally express high levels of glypican-1, or exhibit rapid and specific flow responses as endothelial cells do. For this reason, we can conclude from our experiments only that flow-driven redistribution of GPI-anchored proteins across cell surfaces occurs under physiologically relevant levels of shear stress. Determining whether lateral redistribution of glypican-1 initiates flow mechanosensing and signaling will be the subject of future work.

## Supporting information

Additional details of experimental methods are provided in the supplemental material. In addition, the following supplementary movies are available:

Supplementary Movie 1. Glypican-1-GFP fluorescence in the flat region of the same cell depicted in main text figures 2 and 3 moves downstream immediately after shear stress is applied, while M-SNAP fluorescence remains distributed evenly on the cell surface. The glypican-1-GFP intensity profile reaches a steady state that is maintained as long as the flow is on, as confirmed by the constant exponential fit coefficient.

Supplementary Movie 2. A steady-state gradient of GPI-GFP fluorescence was formed on a COS-7 cell (the same cell depicted in figure 5) under 6.65 Pa of shear stress. After flow stops, the gradient gradually dissipates. We fit the intensity profiles at each time point to cosine functions and determine the diffusion constant by fitting the cosine amplitude time decay.

Supplementary Movie 3. GPI-GFP (left, cyan) undergoes minimal redistribution under low shear stress (2.37 Pa), but a fluorescent antibody to GFP (right, magenta) redistributes strongly toward the downstream edge of the cell.

Supplementary Movie 4. While glypican-1-GFP fluorescence (left, cyan) moves in the downstream direction under shear stress, LifeAct signal (right, magenta) does not.

## Acknowledgments

We thank Daira Santana Almanzar and Maria Jimenez for assistance with buffer viscosity measurements. This research was supported by the United States National Institutes of Health via the National Institute of General Medical Sciences under award R01GM143320 (A.H.S, D.T) and by the Charles E. Kaufman Foundation (A.H.S., D.T).

## Notes

### Competing Interest Statement

The authors have declared no competing interest.

