## supplemental information for "Flow transports extracellular lipid-anchored proteins across the surface of living COS-7 cells"

### Supplemental material for Flow transports GPI-anchored proteins across the surface of living COS-7 cells

**Determining glypican-1-GFP molecular weight.** COS-7 cells were grown in T75 flasks and transfected with glypican-1-GFP pcDNA3.1+ as described in the methods section. The morning following transfection, cells were treated with Trypsin-EDTA, 0.25% (Gibco) for 5 minutes at 37 °C to unadhere from the culture flask. Cells were pelleted at 4000xg for 5 minutes and resuspended in Cell Extraction Buffer (Invitrogen) with 1 mM phenylmethylsulfonyl fluoride (PSMF) and 1X Halt Protease and Phosphatase Inhibitor Cocktail (Thermo Scientific). After resuspension and cell lysis, the sample was pelleted again at 20000xg for 10 minutes. Total protein concentration of the supernatant was determined by BCA Protein Assay Kit (Pierce). For the sample treated with Bacteroides Heparinase III(New England Biolabs, P0737), cell lysate in cell extraction buffer was diluted with water and mixed with 10 $\mu$L 10X Heparinase Reaction Buffer and 1 $\mu$L HepIII enzyme in a final volume of 100 $\mu$L. This reaction was allowed to proceed for 21 hours at 30 °C and then inactivated by heating at 100 °C for 1 minute. An untreated sample was prepared by diluting the same volume of cell lysate with water up to 100 $\mu$L. Each sample (HepIII-treated and untreated) was mixed with 4X SDS loading dye and boiled prior to loading on a 10% acrylamide SDS-PAGE. 15.5 $\mu$g of total protein was loaded per well for the treated and untreated samples; the molecular weight marker used was PageRuler Plus Prestained Protein Ladder (Thermo Scientific). The gel was run at 130 V until the dye front reached the bottom of the gel. Amersham Protran Premium Nitrocellulose Membrane (Cytiva) was soaked in cold transfer buffer (25 mM tris, 192 mM glycine, 20% v/v methanol) along with the gel and extra thick blot filter paper (Bio-Rad) for 5 minutes. The gel was transferred on to the nitrocellulose membrane using a Trans-Blot Turbo Transfer System (Bio-Rad) according to manufacturer instructions. The membrane was blocked with 5% bovine serum albumin (BSA) in 1X TBS-T (0.1% Tween) for 1 h at RT, and then washed three times with 1X TBS-T for 5 minutes each. The anti-GFP primary antibody (Cell Signaling D5.1, # 2956) was used at a 1:1000 dilution in 5% BSA in 1X TBS-T and was incubated overnight at 4 °C. After primary incubation, the membrane was washed three more times with 1X TBS-T for 5 minutes each and incubated with anti-rabbit IgG HRP-linked secondary antibody (Cell Signaling, # 7074) at 1:4000 dilution in 1X TBS for 1 h at RT. The membrane was washed five times with 1X TBS-T for 5 minutes each before adding Clarity Max Western ECL Substrate (Bio-Rad) for imaging.

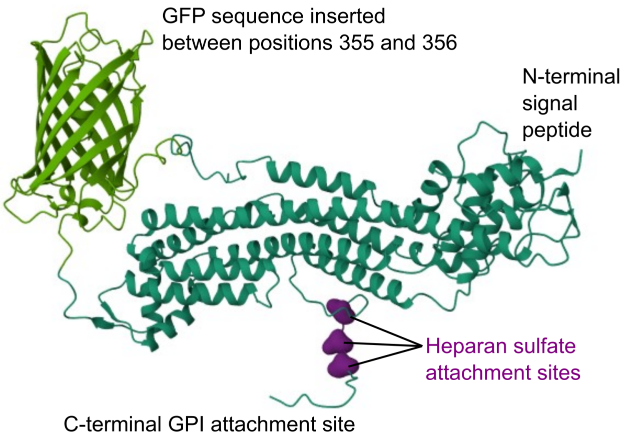


**Fig. 1.** Structure of the glypican-1-GFP construct. We inserted the sequence for EGFP between residues 355 and 356 of the glypican-1 sequence, located on the opposite side of the folded region from the GPI anchor. A mid-sequence insertion was required to avoid interfering with either the GPI anchor or surface localization of the protein. Purple blobs are Gaussian surface volumes highlighting Ser 486, Ser 488 and Ser 490, which are the sites for heparan sulfate attachment. This structure was generated from the plasmid sequence with AlphaFold2 using the COSMIC[2] platform and rendered using Mol* (1).

**Calculation of the Anti-GFP antibody to GPI-GFP molar ratio.** To calculate the number of anti-GFP antibodies bound per GPI-GFP protein, we compared the intensity of GPI-GFP and anti-GFP signals in the cell movies with that from protein solutions of known concentrations. We used a known concentration of anti-GFP antibody solution (Molecular weight: 150 kDa) and monomeric streptavidin-GFP protein solution (Molecular weight: 42.8 kDa) as intensity controls. We calculated the molarity of the solutions from their known concentrations and molecular weights. We captured the images of this solution at the same laser intensity and exposure time settings as those

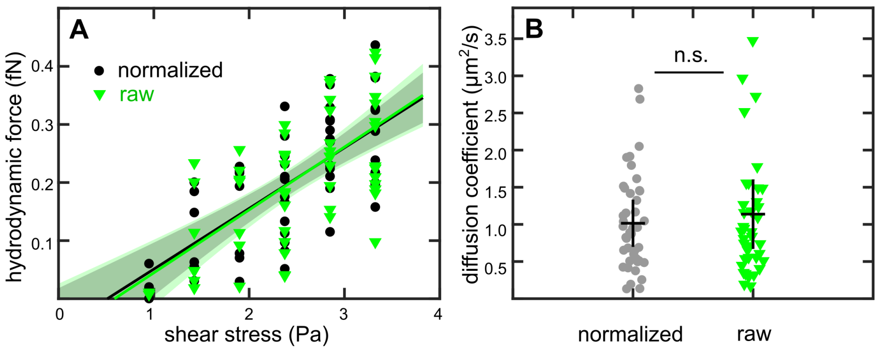


**Fig. 2.** Normalizing by M-SNAP signal did not alter average results. (A) We analyzed glypican-1-GFP fluorescence normalized by fluorescence from M-SNAP (black circles), and then again with no normalization (green triangles). For hydrodynamic force experiments, half of the first-frame fluorescence signal was subtracted in both cases. Although individual hydrodynamic force values changed due to normalization, the average results were not significantly different. The average hydrodynamic force per Pascal of shear stress was 0.104 $\pm$ 0.034 fN for the raw data, and 0.100 $\pm$ 0.027 fN for normalized data. Solid lines show the fitted slope, shaded areas indicate 95% prediction intervals for the fit. In a small number of the non-normalized cases, the exponential fit of raw data failed; those results were not included in the average. Panel (B) shows that there was also no significant difference in the mean diffusion constant determined from raw data (green triangles, 1.14 $\pm$ 0.93 $\mu m^2/s$) and from data normalized by the M-SNAP signal (gray circles, 1.02 $\pm$ 0.64 $\mu m^2/s$). Horizontal lines show the mean value and vertical lines are one standard deviation in length.

used for the antibody cell flow experiment (Figure 8). We drew a region of interest (ROI) for the control and measured mean intensity across the ROI for both mSA-GFP and anti-GFP solutions. This ROI was kept the same for intensity measurement from the one confocal slice of the cell images (in the flat region) from the flow experiment. From the intensity ratio of anti-GFP to mSA-GFP control for a known molar ratio, we determined the unknown molar ratio of anti-GFP to GPI-GFP corresponding to the intensity ratio obtained from the cell flow experiments. The molar ratio of anti-GFP to GPI-GFP proteins varied between 1:80 and 1:300 for the five different cells we calculated. Here, we assume that the solution conditions or GFP fusion protein modifications did not alter the fluorescence of GFP. Nevertheless, we can safely say that not all the GPI-GFP proteins bind with an anti-GFP antibody.

DRAFT

1. M. A. Cianfrocco, M. Wong-Barnum, C. Youn, R. Wagner, and A. Leschziner. COSMIC2: A Science Gateway for Cryo-Electron Microscopy Structure Determination. In *Practice and Experience in Advanced Research Computing 2017: Sustainability, Success and Impact*, PEARC '17, pages 1–5, New York, NY, USA, July 2017. Association for Computing Machinery. ISBN 978-1-4503-5272-7. doi: 10.1145/3093338.3093390.

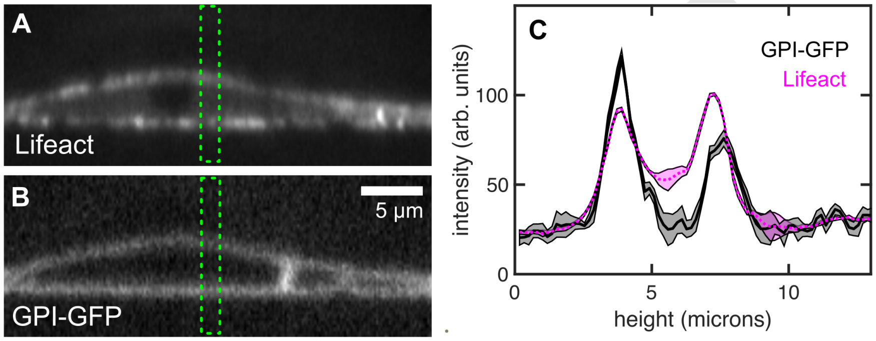


**Fig. 3.** Actin fluorescence signal. Cross-sections of confocal stacks through the same COS-7 cell as depicted in main text figure 8. The cell was doubly transfected with LifeAct (A) and GPI-GFP (B). Panel (C) shows average fluorescence intensity inside the green outlines in panels A and B, which cut through the perinuclear region. GFP fluorescence inside the cell (black solid line) drops to the same level observed outside the cell, indicating that the majority of the GFP fluorescence is located at the membrane. In contrast, the LifeAct fluorescence (magenta dotted line) level inside the cell remains high, likely indicating the presence of unpolymerized actin. To avoid including unpolymerized actin in our analysis, we used maximum intensity projections for the LifeAct signal in figure 8.

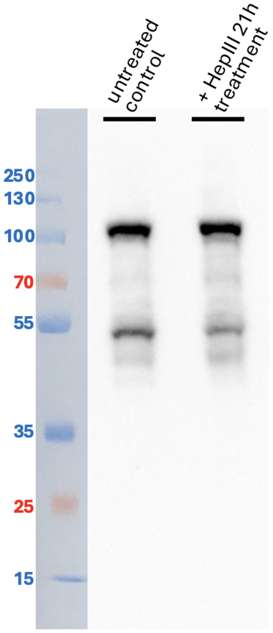


**Fig. 4.** We measured the molecular weight of our glypican-1-GFP construct using a Western blot, detecting an antibody to GFP. From the sequence, the expected molecular weight is approximately 89 kDa; 15.5 $\mu$g of total protein were used per well. Heparinase treatment is expected to collapse the broad signal from high molecular weight heparan sulfate proteoglycan (which is often poorly bound to blotting membranes) down to the core protein with short heparan sulfate stubs, plus N-linked oligosaccharides, a species which is typically also present in heparan sulfate proteoglycan blots of cellular material. The band at 105 kDa likely represents the core protein and N-linked oligosaccharides, but not the full-length heparan sulfate chains. Heparan sulfate chains can be very large (60 kDa per chain in fibroblasts) and are likely to be sheared off of the protein during sample preparation.

DRAFT

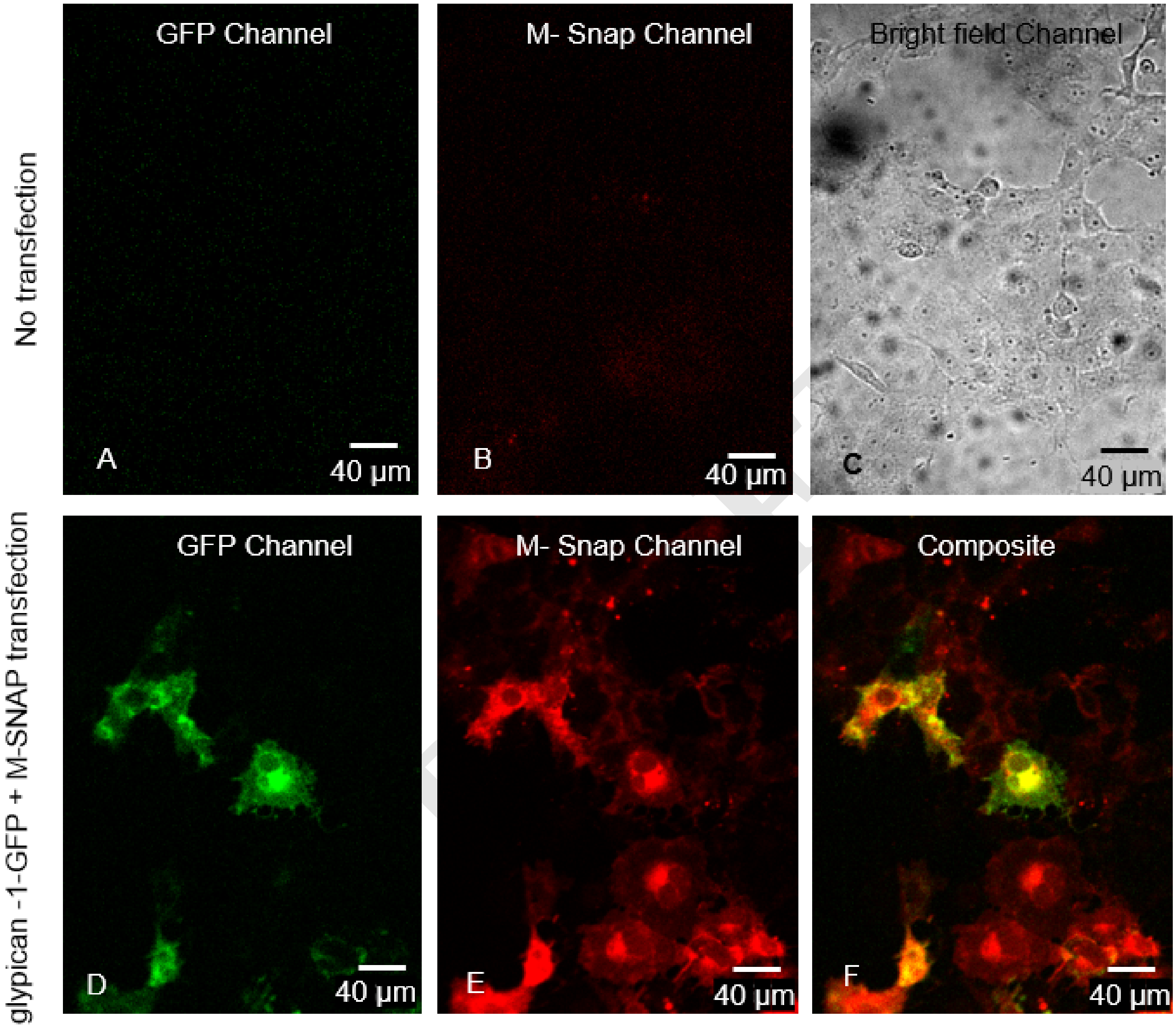


**Fig. 5.** Negative control for glypican-1-GFP and M-SNAP double transfection is shown in panels A and B. The cells were treated with SNAP-Cell TMR-Star substrate in the same way as for transfected cells. A bright-field image showing cells corresponding to panels A and B is shown in panel C. After transfection cells show GFP fluorescence in panel D and SNAP-Cell TMR-Star 580 nm fluorescence in panel E. Panel F shows the composite image of panels D and E.

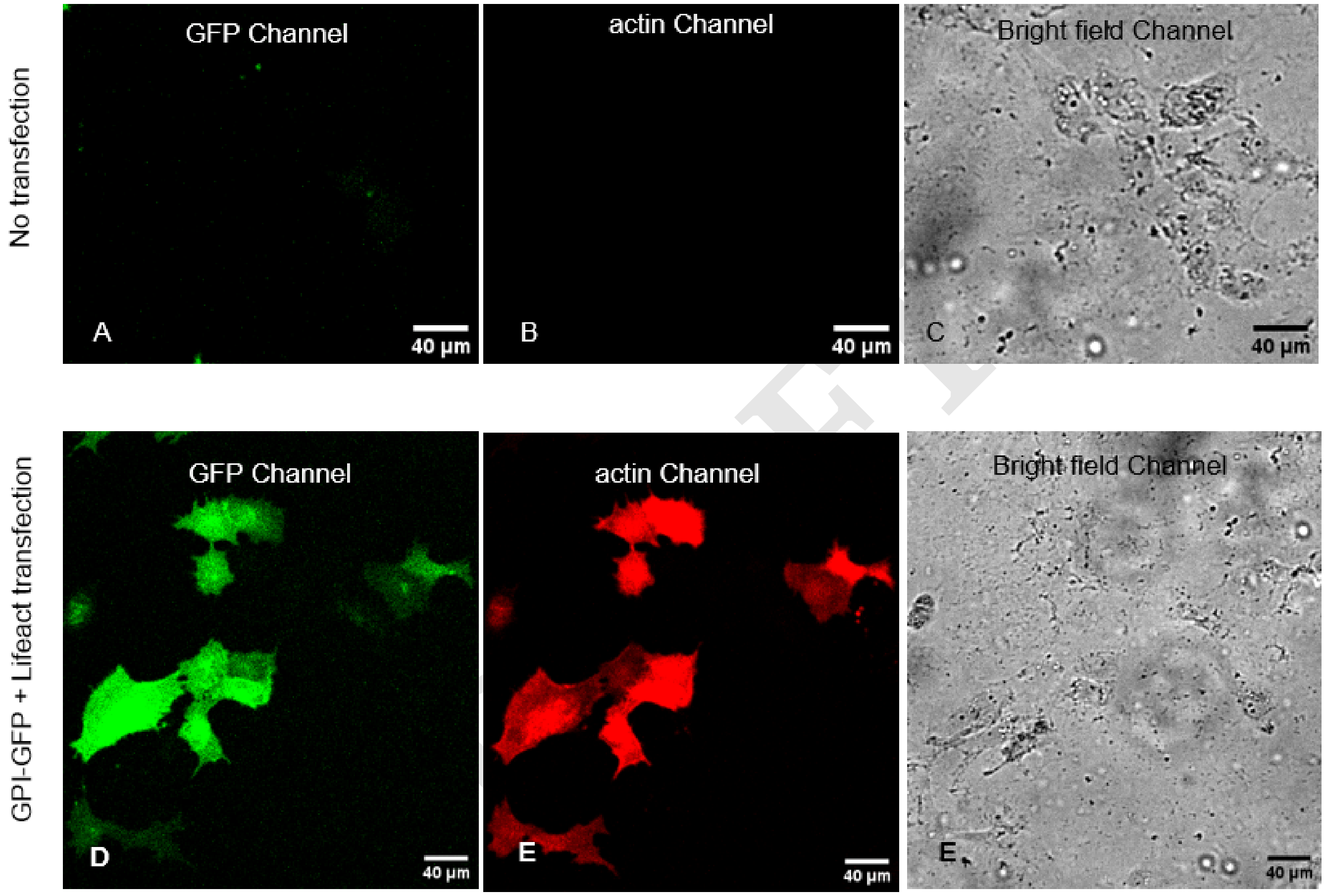


**Fig. 6.** Negative control for GPI-GFP and Lifeact double transfection is shown in panels A and B. A bright-field image showing cells corresponding to panels A and B is shown in panel C. After transfection cells show GFP fluorescence in panel D and Lifeact (red) fluorescence in panel E. Panel F shows the corresponding to bright field image of cells panels D and E.

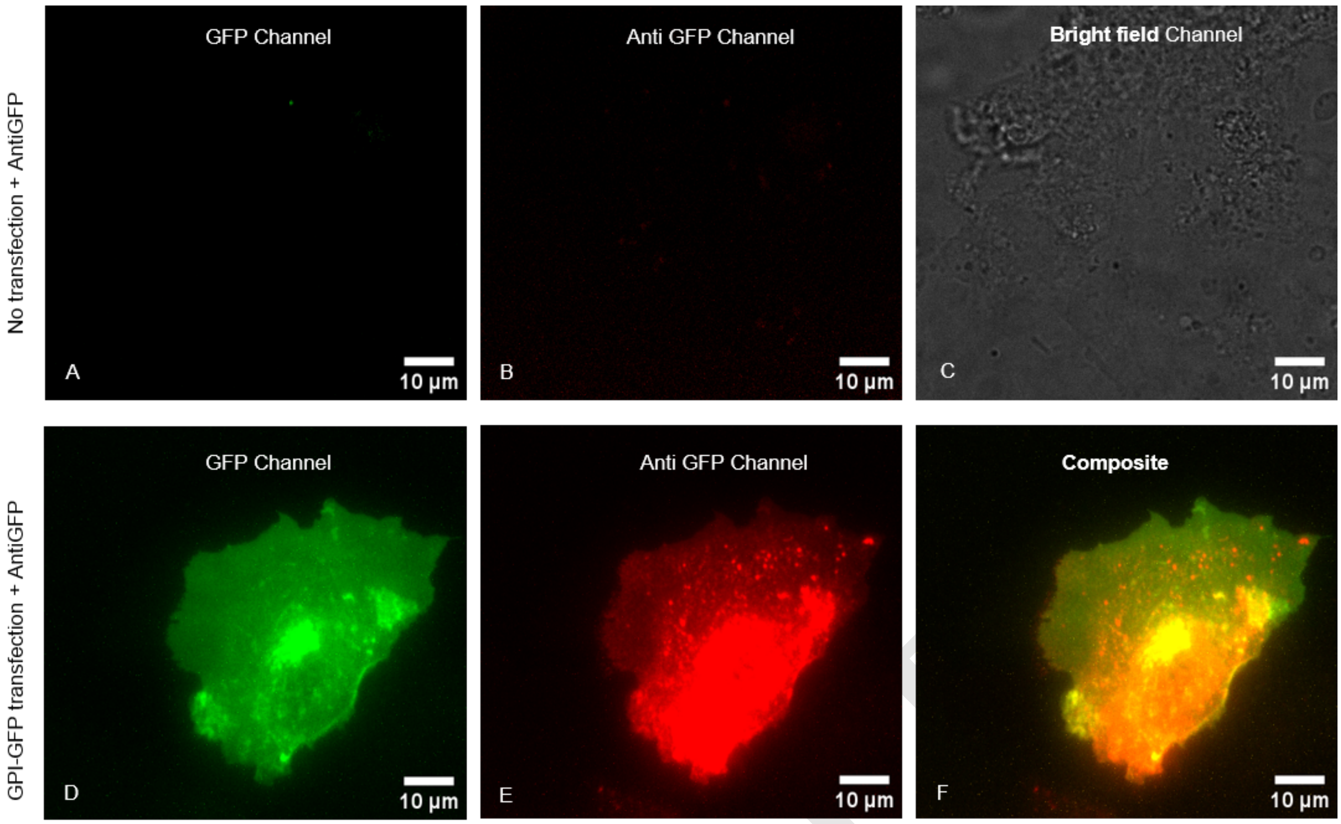


**Fig. 7.** A negative control for anti-GFP labeling is shown in panels A, B, and C. Panel A displays an image of non-transfected cells in the GFP channel. Panel B shows the image of the cells in the anti-GFP channel after treatment with anti-GFP. A bright-field image showing the cell corresponding to panels A and B is shown in panel C. After transfection, cells show GFP fluorescence in panel D, overlapping with specific anti-GFP antibody(red) fluorescence in panel E. Panel F shows the composite image of panels D and E.

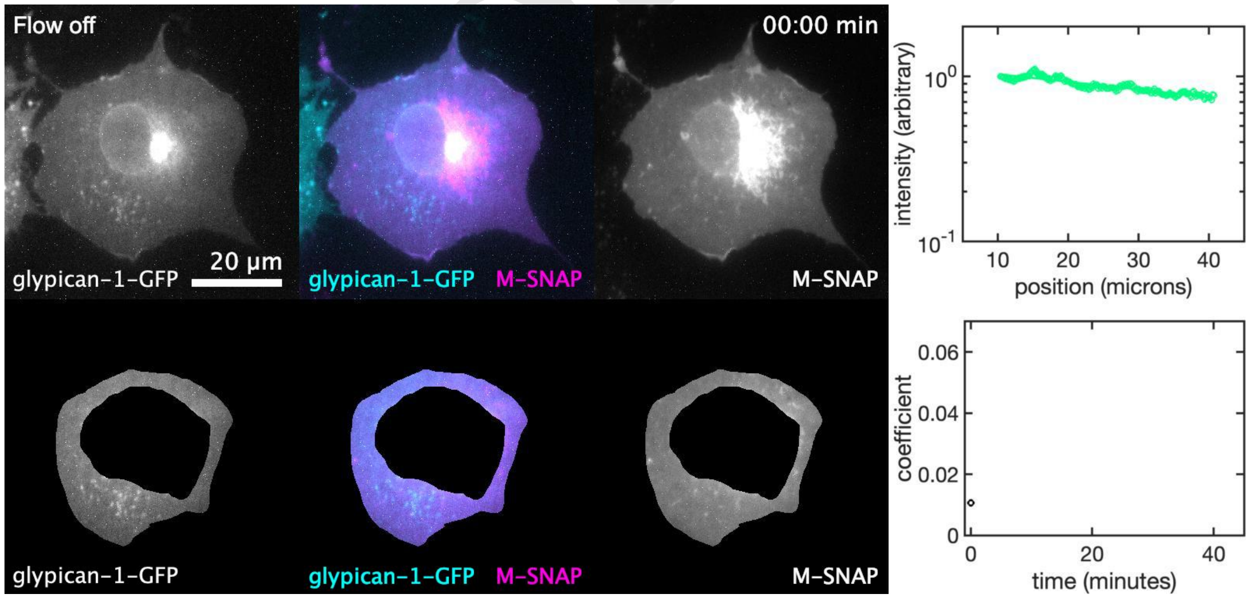


**Fig. 8.** Caption for Movie 1. Glypican-1-GFP fluorescence in the flat region of the same cell depicted in main text figures 2 and 3 moves downstream immediately after shear stress is applied. M-SNAP fluorescence remains distributed evenly on the cell surface. The glypican-1-GFP profile reaches a steady state that is maintained as long as the flow is on, demonstrated by the constant exponential fit coefficient.

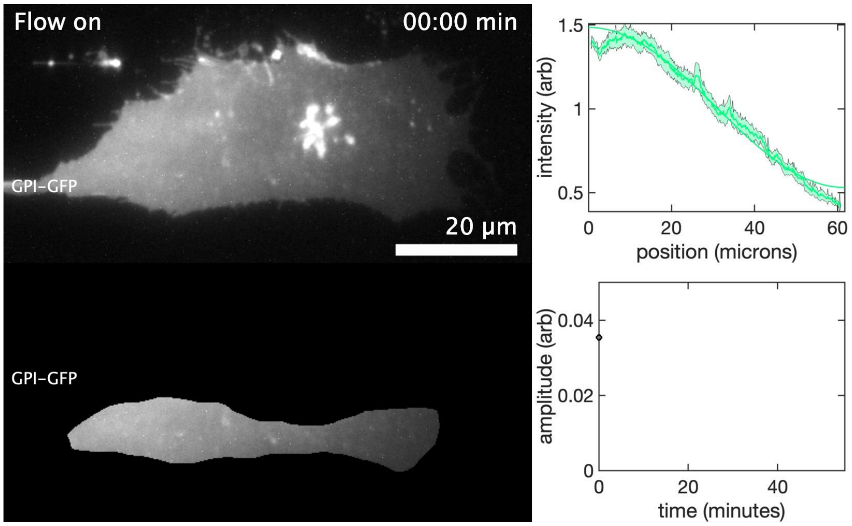


**Fig. 9.** Caption for Movie 2. A steady-state gradient of GPI-GFP fluorescence was formed on a COS-7 cell (the same cell depicted in figure 5) under 6.65 Pa of shear stress. After flow stops, the gradient gradually dissipates. We fit the intensity profiles at each time point to cosine functions and determine the diffusion constant by fitting the cosine amplitude time decay.

DRAFT

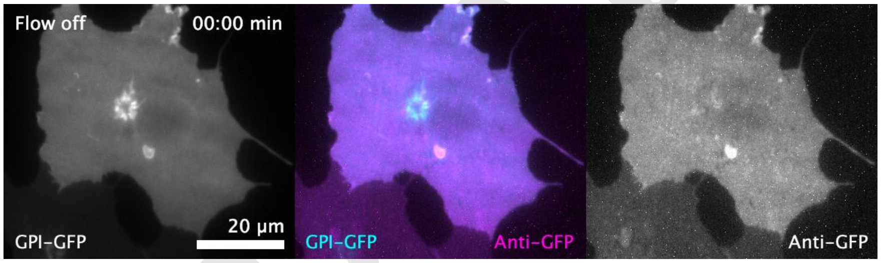


**Fig. 10.** Caption for Movie 3. GPI-GFP (left, cyan) undergoes minimal redistribution under low shear stress (2.37 Pa), but a fluorescent antibody to GFP (right, magenta) redistributes strongly toward the downstream edge of the cell.

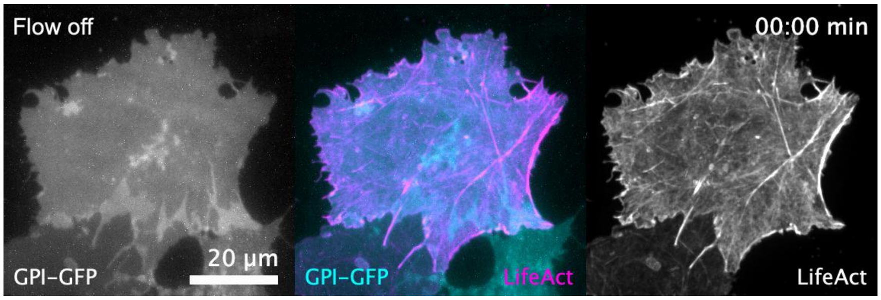


**Fig. 11.** Caption for Movie 4. While glypican-1-GFP fluorescence (left, cyan) moves in the downstream direction under shear stress, LifeAct signal (right, magenta) does not.
